# HPViewer: Sensitive and specific genotyping of human papillomavirus in metagenomic DNA

**DOI:** 10.1101/208926

**Authors:** Yuhan Hao, Liying Yang, Antonio Galvao Neto, Milan R. Amin, Dervla Kelly, Stuart M. Brown, Ryan C. Branski, Zhiheng Pei

## Abstract

**Background:** Shotgun DNA sequencing provides sensitive detection of all 182 HPV types in tissue and body fluid. However, existing computational methods either produce false positives misidentifying HPV types due to shared sequences among HPV, human, and prokaryotes, or produce false negative since they identify HPV by assembled contigs requiring large abundant of HPV reads.

**Results:** We show that HPV shares extensive simple repeats with human and prokaryotes and homologous sequences among different HPV types. The shared sequences caused errors in HPV genotyping and the repeats of human origin caused false positives in HPVDetector. Programs, such as VirusTAP and Vipie, which require *de novo* assembly of shotgun reads into contigs, eliminated false positives at a cost of substantial reduction in sensitivity. Here, we designed HPViewer with two custom HPV reference databases masking simple repeats and homology sequences respectively and one homology distance matrix to hybridize these two databases. It directly identified HPV from short DNA reads rather than assembled contigs. Using 100,100 simulated samples, we revealed that HPViewer was robust for samples containing either high or low number of HPV reads. Using 12 shotgun sequencing samples from respiratory papillomatosis, HPViewer was equal to VirusTAP, and Vipie and better than HPVDetector with the respect to specificity and was the most sensitive method in the detection of HPV types 6 and 11. We demonstrated that contigs-based approaches had disadvantages of detection of HPV. In 1,573 sets of metagenomic data from 18 human body sites, HPViewer identified 104 types of HPV in a body-site associated pattern and 89 types of HPV co-occurring in one sample with other types of HPV at least once.

**Conclusions:** We demonstrated HPViewer was sensitive and specific for HPV detection in metagenomic data. It was also suggested that masking shared sequences is an effective approach to avoid false positive detection and identifying HPV from short metagenomic reads is more sensitive than assembled contigs. The innovative homology distance matrix connecting two HPV databases, repeat-mask and homology-mask, optimized the balance of sensitivity and specificity. HPViewer can be accessed at https://github.com/yuhanH/HPViewer/.

## Background

Human papillomavirus (HPV) is a type of double-stranded small DNA virus that causes nearly 610,000 cases of cancers annually in the world [1]. Currently, 210 types of HPV have been identified in the International HPV Reference Center (http://www.hpvcenter.se) and this number is increasing monthly. There are 182 types of HPV with complete genomes sequences in the PapillomaVirus Episteme (PaVE) (https://pave.niaid.nih.gov/). Many studies have demonstrated that HPV is a vital cause of cervical cancers [2–4] and these studies have classified HPV types as high risk and low risk. Munoz et al., grouped HPV 16, 18, 31, 33, 35, 39, 45, 51, 52, 56, 58, 59, 68, 73, 82 as high risk; and HPV 6, 11, 40, 42, 43, 44, 54, 61, 70, 72, 81, 89 were considered as low risk [5]. HPV has also been linked to other cancers including cancers of the oropharynx [6], head, neck [7–9]. Of particular concern, the incidence of HPV-associated oropharynx cancer [10] is growing very rapidly. Furthermore, HPV DNA has been detected in cancers of the lung, colon, esophagus, and urinary bladder [11–14].

The traditional clinical HPV detection methods can be classified into three groups: nucleic acid-hybridization assays, nucleic-acid amplification, and antibody-based assays [15]. Nucleic acid-hybridization assays make use of in situ hybridization, which can detect the 13 most high-risk HPV genotypes, including types 16, 18, 31, 33, 35, 39, 45, 51, 52, 56, 58, 59, and 68 through a biotinylated-probe cocktail (GenPoint HPV Probe Cocktail, Dako) or other HPV types with custom designed probes [16]. Inno-LiPA [17] can detect 32 types of HPV by PCR amplification of a 65-bp region of the conserved L1 gene and then performing reverse line blot hybridization to identify specific HPV types. A real-time TaqMan PCR assay can also be used for HPV detection through determining the presence of mRNA of E6 genes of HPV [18]. There are two FDA-approved HPV assays using nucleic-acid amplification. Cobas® HPV Test by Roche (Indianapolis, IN, USA) can detect 14 types of high-risk HPV DNA through PCR and fluorescence [19]. Aptima® by GenProbe (Woburn, MA, USA) targets high-risk HPV mRNA from E6/E7 genes by transcription-mediated amplification [20]. An indirect assay for HPV16 infection is available by immunohistochemistry of expression of a human gene, p16, because there is an overexpression of p16 resulting from HPV-16 integration into the host genome and disruption of the retinoblastoma pathway [21]. More recently, Lavezzo et al. (2016) proposed a new HPV genotyping method depending on conserved PCR primers for the E6/E7 region [22], but this new method is limited to detection of only high-risk types of HPV.

These methods, although covering mainly the 26 high/low risk HPV types, are sufficient to detect all HPV types related to cervical cancer [23]. Our understanding of the causality of HPV in other cancers is mainly derived from surveys by using the cervical HPV detection methods. However, HPV type distribution in cervical cancers among women from different populations have heterogeneity [24] and there have been no methods or kits specially designed for detecting HPV types found in oropharynx cancers. Thus, HPV prevalence in these cancers could be underestimated due the inabilities of cervical HPV kit to detect all HPV types, so a broad range method to detect all HPV types is needed to allow a complete evaluation of the role of HPV in cancers outside of the uterine cervix.

Shotgun sequencing of human tissue samples or body fluids is a robust tool which can broaden the narrow spectrum of the traditional HPV detection approaches. It depends on bioinformatics pipelines to identify and genotype HPV reads from a large pool of human and microbial DNA sequences. Johannsson et al., (2013) applied MEGABLAST to filter out human and bacteria reads and performed de novo assembly to obtain long contigs and used BLASTn against GenBank to identify HPV [25]. Ma et al., (2014) applied a HPV genotyping framework through BLAST to a local reference HPV database for detection of HPV reads in datasets generated from a variety of human body sites by whole genome shotgun sequencing (WGS) [26]. BLAST is a powerful but time-consuming tool [27] and it is very inefficient for processing millions of short DNA fragments from metagenomic data. HPVDetector, developed in 2015 [28], depends on the Burrows-Wheeler Aligner [29] to match shotgun reads to their reference genome database. There are also several software programs designed for identifying all viruses including HPV in whole genome shotgun sequencing (WGS) data, such as Metavir2 [30], VirSorter [31], VirusTAP [32], VirusScan [33], Vipie [34], VIP [35], and VirFinder [36]. Table 1 provides a summary of important characteristics of 9 different programs available.

**Table 1.**
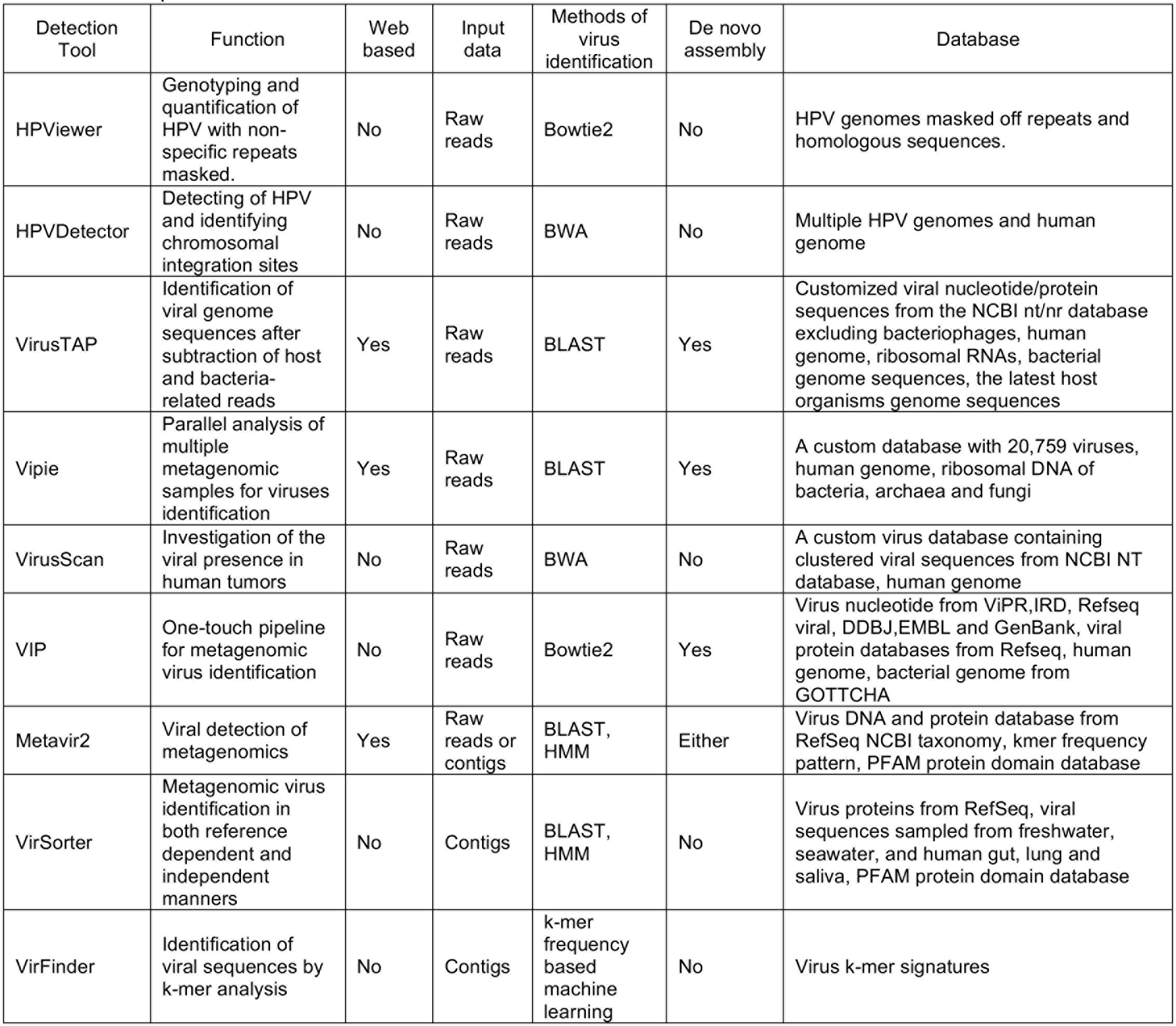
Comparison of current HPV or virome detection tools

One consideration for identifying HPV with short reads in WGS data is false positivity caused by homologous sequences and/or repeats shared among the host, microbes, and HPV. In addition, genotyping can be inaccurate when HPV reads detected are shared by more than one HPV type. One approach to reduce false positivity is using *de novo* assembly to generate large contigs that cover a larger region of HPV genome beyond the shared region. Of the 9 programs, 6 applied or required the *de novo* assembly approach, including VirSorter, VirFinder, Metavir2, VirusTAP, Vipie, and VIP. However, contigs from *de novo* assembly can be constructed only if the data have sufficient coverage, limiting its capability of detecting HPV in samples in which HPV reads are too few to form a contig. Another approach to reduce false positivity is to filter out host and bacterial genome sequences. For example, VIP VirusTAP, and VirusScan subtract the input DNA fragments which can align to the host genome before searching for HPV DNA. This strategy has two shortcomings because of the large size of host genome. It not only takes long time to align input DNA fragments to the host genome but also needs large storage space for the host genome database for local use. In addition, this approach does not reduce genotyping errors due to homology among closely related HPV genotypes. HPVDetector, the program specially designed for HPV detection, does not consider the false positive issues from the host genome and homology among different HPV types.

In the present study, we developed a new HPV detection program – HPViewer that reduces false detection of HPV DNA by masking simple repeats commonly shared among the human genome, prokaryotes, and homologous sequences shared by different HPV types. We evaluated the sensitivity and specificity of HPViewer using 100,100 simulation samples, and in a WGS dataset from patients with recurrent respiratory papillomatosis which are known to be associated with HPV6/11 [37], compared the performance of HPViewer with HPVDetector, VirusTAP and Vipie. We also applied HPViewer to define HPV prevalence distribution and explore the co-occurrence pattern of HPV types in different body sites of healthy samples from the Human Microbiome Project (HMP).

## Results

### HPV genomes share simple repeats with human and prokaryotic genomes

Metagenomic data usually consists of fragments of human and prokaryotic genomes. To detect human and prokaryotic DNA sequences that may interfere with HPV identification, we compared 182 HPV genomes [38] with human genome (GRCh38) using BLASTn [27] and found 165,118 matches (identity >90%, alignment length >50 bp) between 14 HPV types and all human chromosomes (Fig 1 a,b). All matches were simple repeats and most were TA (83.94%) and TG (15.94%) repeats. Other less abundant repeats, such as TTC, TTCTCC and CATA were also found. In particular, low risk pathogenic HPV types 6, 72, 73 share simple sequence repeats with human chr3, chr1, chr1 and chrY, respectively.

**Figure 1.**
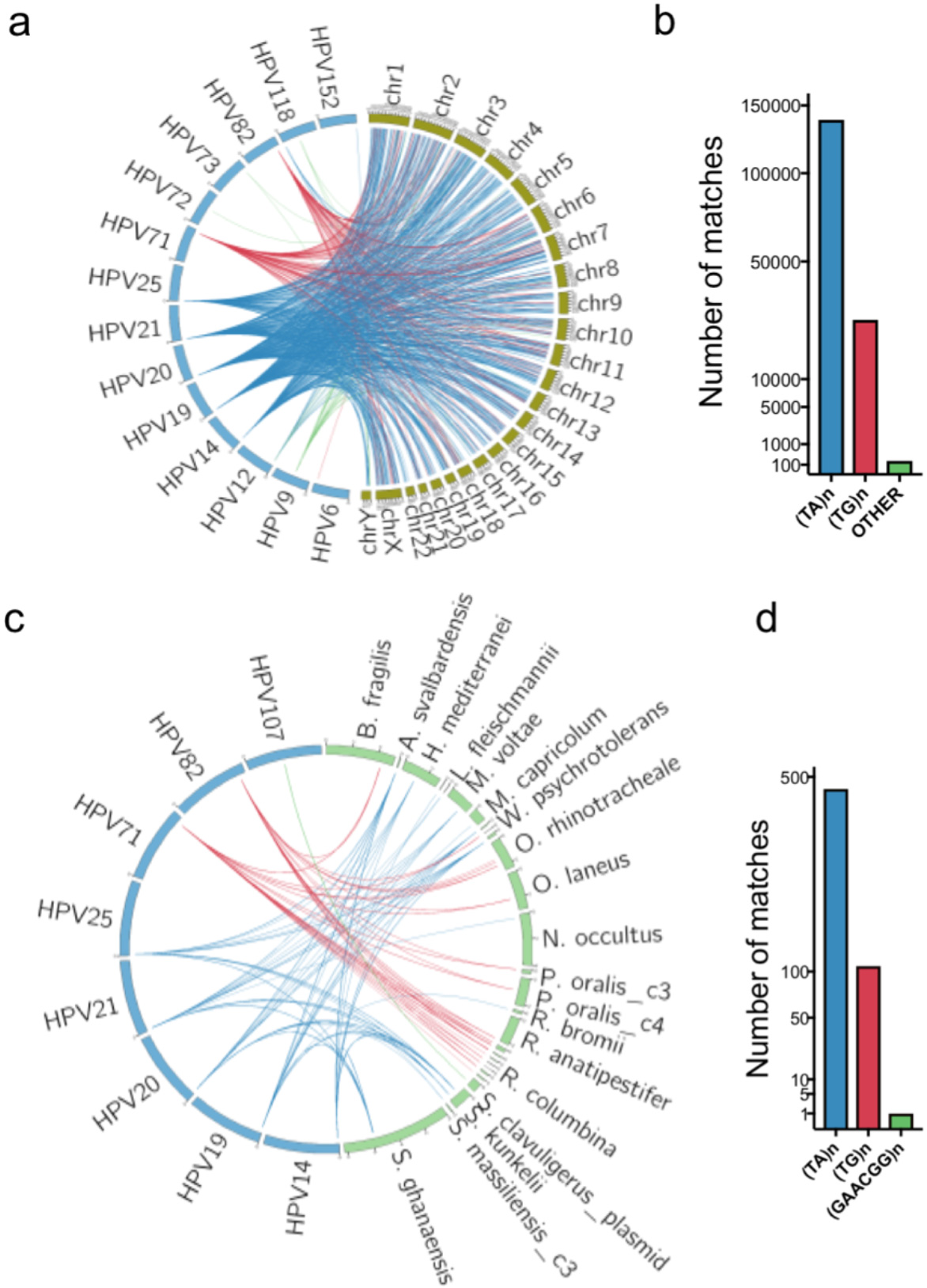
The HPV shared sequences between human and prokaryotic genomes. (**a,b**) The shared sequences between HPV and the human genome. Each line represents one BLASTn alignment. The blue lines represent TG repeats, and the red lines represent TA repeats and green lines represent other repeats alignments. (**c,d**) The shared sequences between HPV and prokaryotic genomes. Most were TG and TA repeats. The only other type shared repeat was GAACGG repeat between HPV107 and *Streptomyces clavuligerus* Plasmid pSCL2 (NZ_CP016560.1). The mapped prokaryotic genomes were listed in **Supplementary Table 1.**

Using the same strategy, we also compared HPV genomes with 1,781 prokaryotic genomes (NCBI 112 prokaryotic reference genomes and 1,669 NCBI prokaryotic representative genomes) (**Supplementary Table 2**) and found 575 matches between 8 HPV types and 18 prokaryotic species (Fig 1 c,d), mainly TA (81.22%) and TG (18.61%) repeats plus GAACGG repeats (0.17%). None of the 8 HPV types were high or low risk cervical HPV types. In all 1,375,680 bp of the 182 HPV genomes, simple repeats accounted for 16,359 bp (1.19%).

### Homologous sequences are widely shared among HPV types

Besides homology between HPV and other organisms, homology among HPV types could also interfere with HPV genotyping. To estimate the extent of homology, we aligned each complete HPV genome with genomes of all other type by sliding all possible 100-bp DNA fragments along its entire genome, with a 90% identity threshold. The degree of homology between different types of HPV varied greatly. There are 29 HPV types which lacked homology with any other HPV type, but 85.9% of HPV76 genome was homologous with other HPV types. In the 182 HPV genomes, 368,789 bp (26.81%) were homologous between two or more HPV types.

### Design of HPViewer

We took a novel, masking approach to minimize the impact of the shared sequences on HPV genotyping. Instead of filtering shared sequences by alignment of millions of raw reads in each sample to human and prokaryotic genomes, we masked the simple repeat sequences in the reference HPV genome database with RepeatMasker [39]. We then compared these masked HPV genomes with human and prokaryotic genomes and found no matches, indicating that our repeat-mask strategy eliminated false positive calling of human or prokaryotic DNA reads as HPV. Next, we masked all homologous regions shared among HPV types as well as simple repeats as our homology-mask strategy. We found the repeat-mask removed only a few hundreds of nucleotides, while homology-mask considerably changed the distribution of HPV effective genome lengths (Fig 2). Finally, we built a homology distance matrix and a homology tree only using homologous sequences shared by any other type of HPV (**Supplementary Fig 1**).

**Figure 2.**
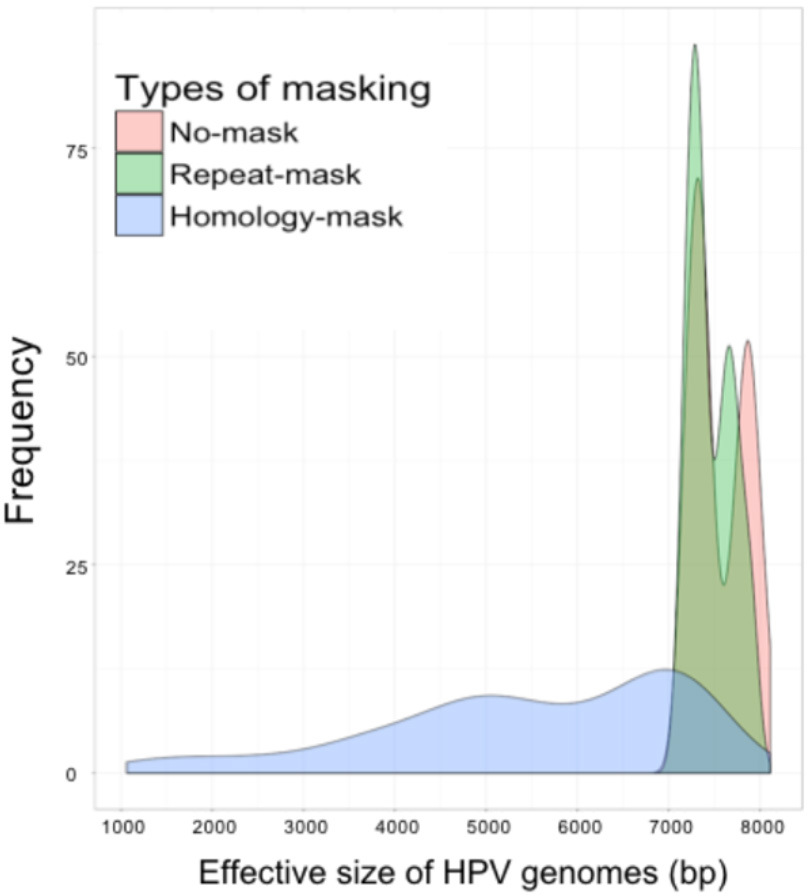
Distribution of HPV effective genome sizes among original HPV genomes, and HPV genomes in repeat-mask, and homology-mask databases. For our mask strategies, the length of HPV genomes was not changed and we called non-N sequences of HPV genome as the effective genome. In sum, the HPV effective genome length ranged from 7,100-8,104 bp for original genomes, 7,100-7,995 bp for repeat-mask genomes, and 1,061-7,698 bp for homology-mask genomes.

### Estimation of sensitivity and specificity of HPViewer using simulated data

We developed HPViewer for specific detection and quantification of HPV from metagenomic data. Initially, we planned to use a repeat-mask mode to eliminate false positivity caused by human and prokaryotic genomes and a homology-mask mode to prevent errors in genotyping among closely related HPV types.

We evaluated these two modes with 100,100 simulated samples composed of 143 HPV types at various sequencing depths. The sensitivity progressively increased with higher sequence depth for both modes. At the depth of 2 reads/sample, the repeat-mask mode (76.8%) was much more sensitive than the homology-mask mode (29.4%) due to overlooking true HPV reads shared among different HPV types by the homology-mask mode (Fig 3a). Sensitivity reached a plateau (>98.9%) at 50 reads for both modes. The specificity was ~100% for both modes at 2-10 reads and was maintained for the homology-mask mode up to 1,000 reads. However, the specificity progressively decreased at >50 reads and dropped to 89.8% at 1,000 reads for the repeat-mask mode due to errors in genotyping of closely related HPV types (Fig 3b). In contrast, the currently available software HPVDetector [28] was less specific than both modes and less sensitive than the repeat-masked mode (Fig 3).

**Figure 3.**
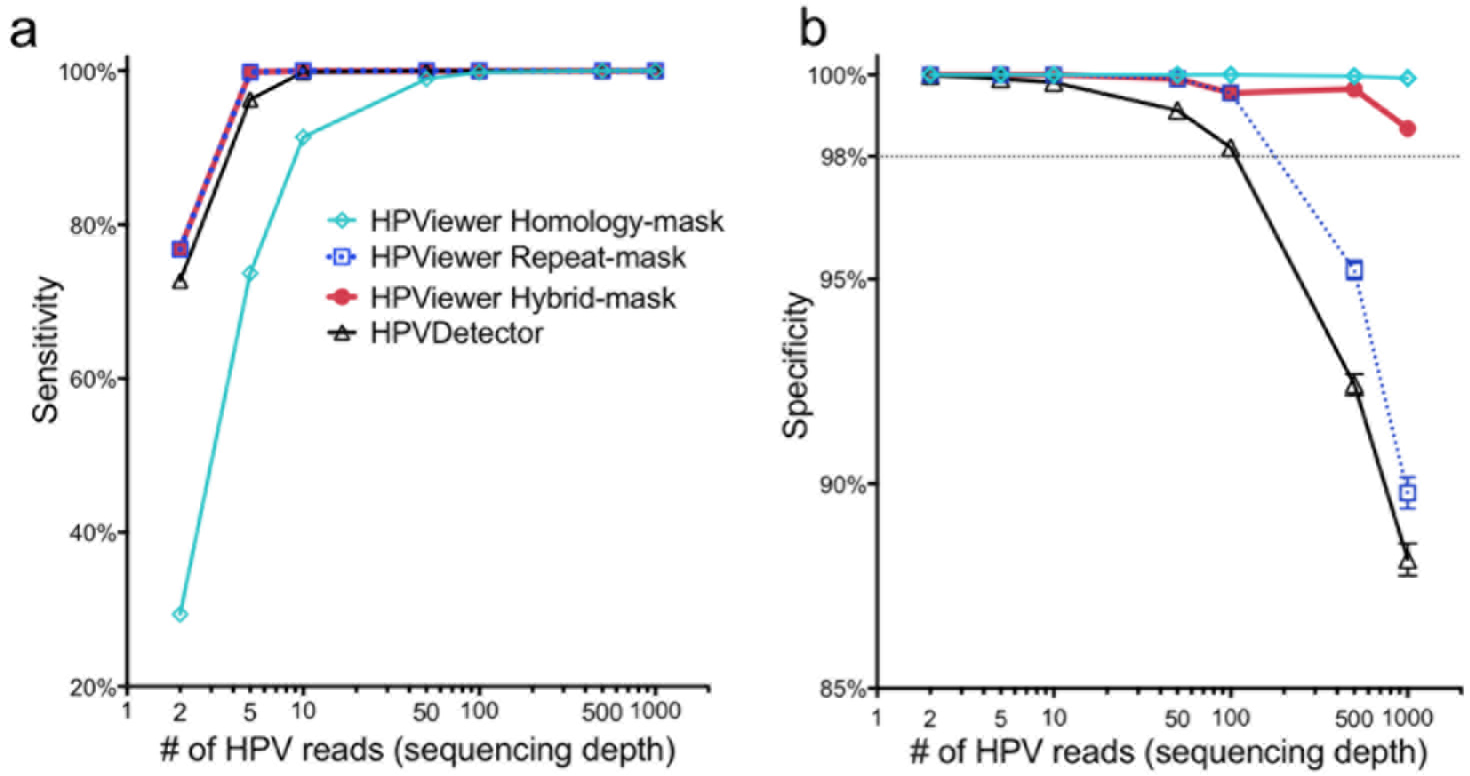
Evaluation of HPViewer using simulated HPV shotgun sequencing data. We compared the performance of three modes of HPViewer and HPVDetector using 100,100 simulated HPV samples. (**a,b**) Comparison sensitivity and specificity of HPViewer three modes and HPVDetector using simulated single HPV data with different sequencing depths: 2, 5, 10, 50, 100, 500, 1000 reads. Under each sequencing depth, we simulated each HPV type 100 samples. Considering HPVDetector only contained 143 types of HPV genomes, we only generated simulated reads from those HPV genomes.

To surmount the low sensitivity of the homology-mask mode and the low specificity of the repeat-mask mode, we created a novel hybrid approach by combining the two modes using the pair-wise homology distance matrix. In this approach, the repeat-mask mode was used first to screen all HPV reads in a sample. If only a single type HPV was detected, the reads were considered as true positive. If multiple HPV types were detected, their homology distance was determined using the pair-wise homology distance matrix. A HPV match with no close relatives (homology distance < 0.35) was counted as true positive while closely related HPV types were examined with the homology-mask mode. Only HPV types re-detected using the homology-mask mode were considered as true positive (Fig 4). The hybrid-mask mode had the same sensitivity of repeat-mask mode, 76.8%, at the depth of 2 reads and improved the specificity of the repeat-mask mode to 98.7% from 89.8% at 1,000 reads (Fig 3). These findings suggest that this hybrid-mask is optimal for detection of HPV in samples that contain either high or low number of HPV reads. This hybrid screening method was set as the default in the distributed version of HPViewer software.

**Figure 4.**
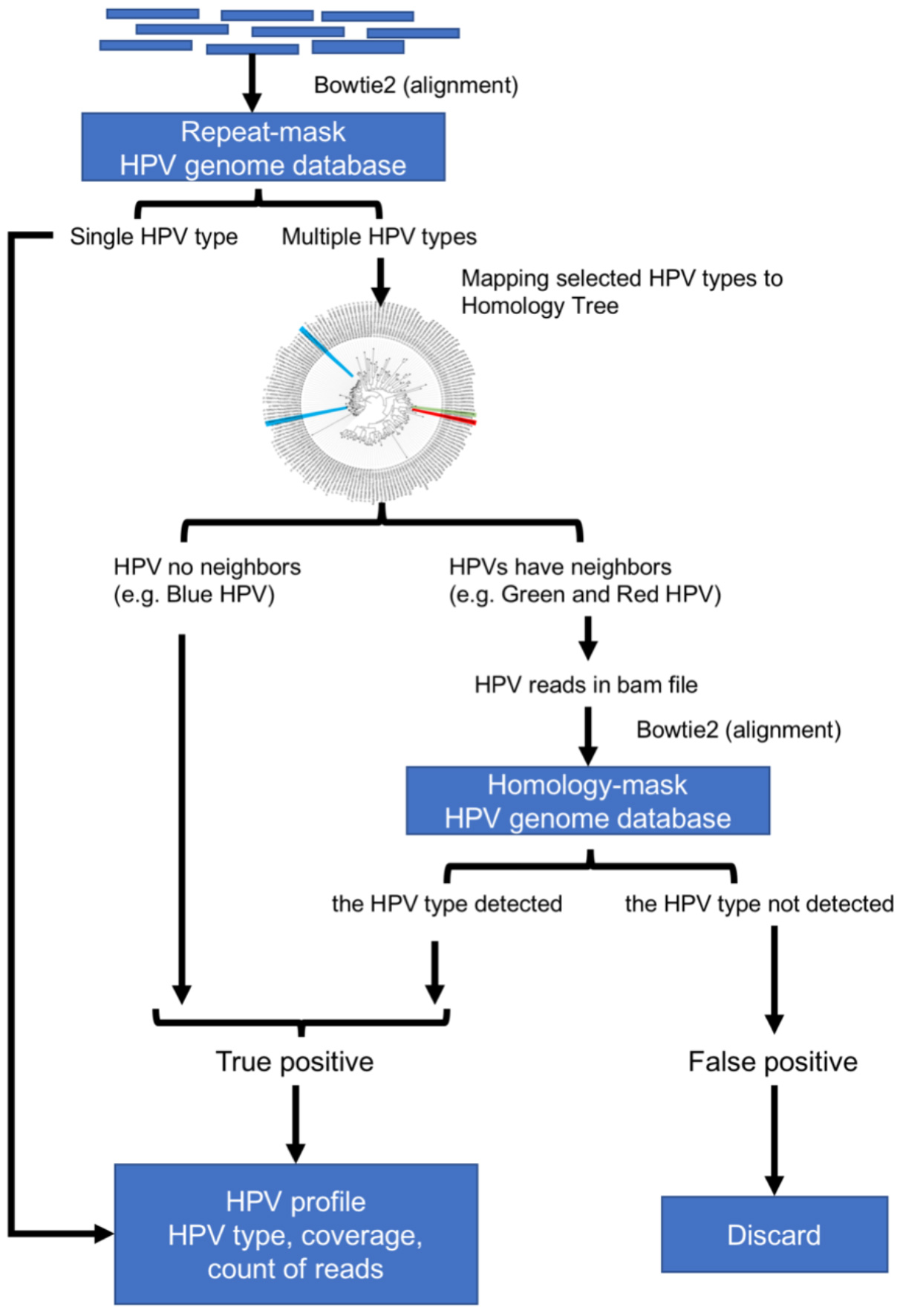
The workflow of hybrid-mode of HPViewer. The hybrid mode of HPViewer is a combination of repeat-mask database and homology-mask database through the homology distance matrix. The input is trimmed fastq file and the output is a table containing HPV types and abundance.

### Comparisons of the performance of HPViewer with HPVDetector, VirusTAP, and Vipie using shotgun sequencing data of recurrent respiratory papillomatosis

We evaluated HPViewer with specimens infected by known HPV types. We performed shotgun sequencing on tumor tissues and matched oral washes from six patients with recurrent respiratory papillomatosis, known to be caused by HPV6 or 11. HPViewer detected HPV6 in 4 tumor tissues and 2 matched oral wash samples and HPV11 in 2 tumor tissues. In contrast, HPVDetector, a standalone program designed to directly genotype HPV reads from raw shotgun sequences, misidentified repeat reads from the human genome as false positive HPV19 (n=4 samples), HPV71 (n=9), or HPV82 (n=7) (Table 2). HPVDetector also misidentified two reads as HPV 11 in sample 7T which matched perfectly to HPV6. For these 12 samples, HPVDetector predicted an average of 1.9 wrong HPV types per sample (Fig 5a). In the tumor tissues, HPVDetector consistently underestimated HPV read counts compared to HPViewer (p=0.028, two-tailed paired t-test), for both HPV6 and HPV11 (Table 2).

**Table 2.** Comparison of HPViewer and other programs on detection and genotyping of HPV in recurrent respiratory papillomatosis

| Sample ID | HPV type: number of reads detected |  |  |  |  |
| --- | --- | --- | --- | --- | --- |
|  | bowtie2 Complete HPV genomes | HPVViewer | HPVDetector | VirusTAP | Vipie |
| 3T | HPV6: 2851<br>HPV71: 1 | HPV6: 2721 | HPV6: 2150 | HPV6 | HPV6 |
| 7T | HPV6: 386<br>HPV71: 1 | HPV6: 361 | HPV6: 260<br>HPV11: 2<br>HPV19: 1<br>HPV71: 1 | No hits | HPV6 |
| 8T | HPV6: 1250 | HPV6: 1194 | HPV6: 929<br>HPV71: 1 | HPV6 | HPV6 |
| 9T | HPV11: 4514 | HPV11: 4229 | HPV11: 3481<br>HPV71: 5 | HPV11 | HPV11 |
| 12T | HPV11: 243<br>HPV71: 1 | HPV11: 228 | HPV11: 167<br>HPV82: 3 | No hits | HPV11 |
| 15T | HPV6: 1285 | HPV6: 1223 | HPV6: 954<br>HPV71: 1<br>HPV82: 1 | HPV6 | HPV6 |
| 3W | HPV6: 2<br>HPV51: 6<br>HPV11: 1<br>HPV71: 7 | HPV6: 2<br>HPV51: 6 | HPV6: 1<br>HPV51: 5<br>HPV71: 7<br>HPV82: 2 | No hits | No hits |
| 7W | HPV6: 2<br>HPV71: 11<br>HPV82: 1 | HPV6: 2 | HPV19: 1<br>HPV71: 3<br>HPV82: 2 | No hits | No hits |
| 8W | HPV19: 1<br>HPV71: 2 | No hits | HPV19: 2<br>HPV71: 3 | No hits | No hits |
| 9W | HPV71: 13<br>HPV82: 2<br>HPV20: 1 | No hits | HPV71: 3<br>HPV82: 5 | No hits | No hits |
| 12W | HPV71: 8<br>HPV82: 1 | No hits | HPV71: 2<br>HPV82: 4 | No hits | No hits |
| 15W | HPV71: 19<br>HPV82: 3 | No hits | HPV19: 1<br>HPV71: 5<br>HPV82: 1 | No hits | No hits |

False positive results are highlighted in red and false negative results in blue.

**Figure 5.**
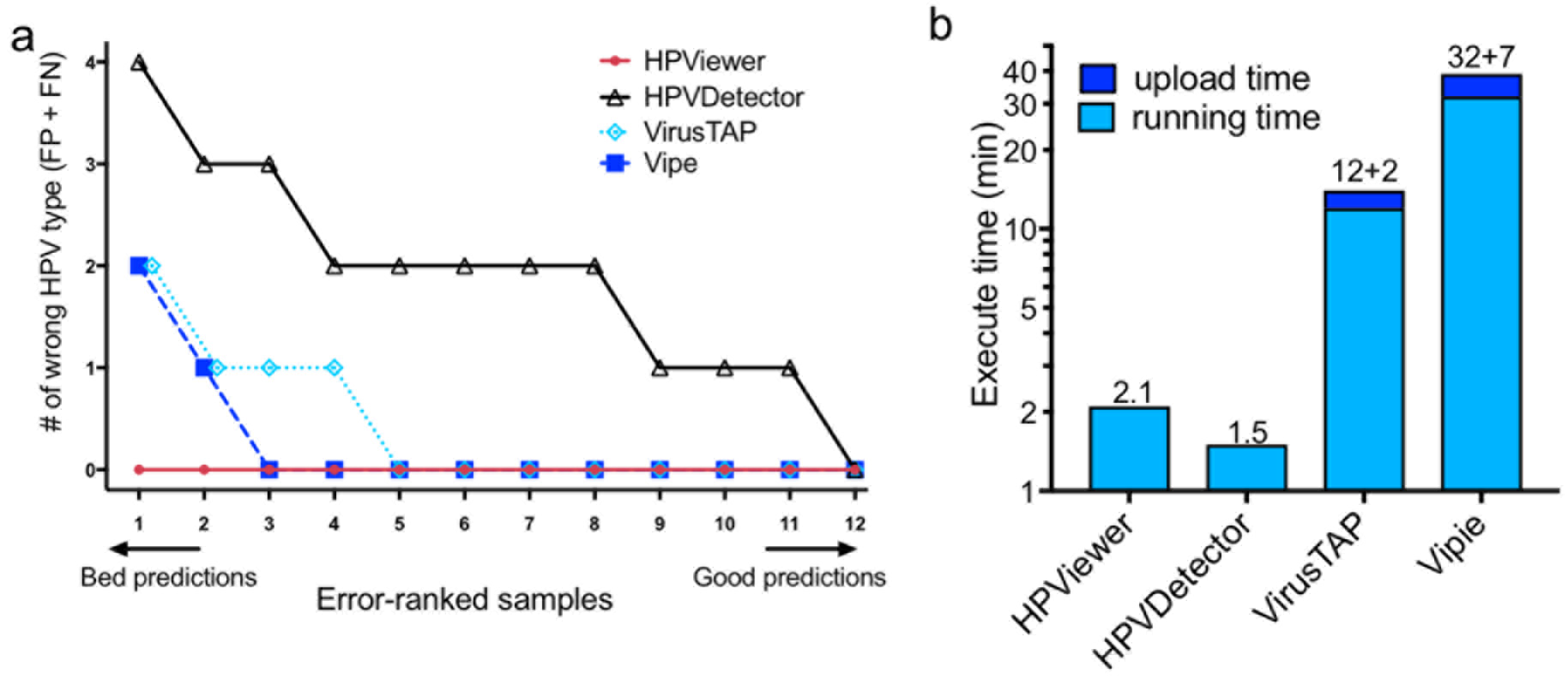
Comparison of different tools on recurrent respiratory papillomatosis shotgun sequencing samples. (**a**) Number of wrong predicted HPV types with respect to 12 shotgun sequencing samples. Wrong predicted HPV types consisted of false positive and false negative types. **(b)** Comparisons of execute time between HPViewer, HPVDetector, VirusTAP, and Vipie for a fastq file containing 7.6 M reads. VirusTAP and Vipie were web-based tools, so they also had a upload time.

VirusTAP is a web-based tool [32] that filters human and bacterial reads and utlizes *de novo* assembly of filtered reads into contigs. It could only detect HPV from samples with very large number of HPV reads. For example, it was able to detect HPV6 in sample 15T in which HPViewer identified 1,223 HPV reads but failed to detect HPV6 in samples 7T and 12T in which HPViewer identified 361 and 228 HPV reads. Vipie is another virus detection program [34] that utilizes *de novo* assembly of all reads into contigs. It was more sensitive than VirusTAP and had equivalent performance with HPViewer in tumor samples with a >100 HPV reads. It successfully detected HPV6 in four tumor samples and HPV11 in two tumor samples. However, it failed to detect HPV in the oral wash samples that contained only a very small number of HPV reads. For example, the 6 HPV51 reads and 2 HPV6 reads in sample 3W and the 2 HPV6 reads in sample 7W were not observed by Vipie.

We compared the computing time of HPViewer with HPVDetector, VirusTAP, and Vipie on analysis of a pair-end fastq file of sample 3T (fastq.gz file, 340 MB, 7.6M reads). The task took approximately two minutes for HPViewer and HPVDetector, 12 minutes for VirusTAP (plus 2 minutes uploading time), and 32 minutes for Vipie (plus 7 minutes uploading time) to complete (Fig 5b). VirusTAP and Vipie cost longer time than HPViewer and HPVDetector to complete the same task because they needed extra time for the process of *de novo* assembly. VirusTAP pre-selects virus reads before the *de novo* assembly on a small number of selected sequences while Vipie performs *de novo* assembly on all reads before identifying HPV contigs. The longer time that Vipie needed than VirusTAP to analyze sample 3T reflects the fact that its scale of *de novo* assembly was much larger than that of VirusTAP.

### Evaluation of HPViewer with shotgun sequencing data from healthy human subjects in the Human Microbiome Project

To evaluate the performance of HPViewer with datasets with unknown HPV status, we downloaded HMP Illumina metagenomic datasets that were originally generated from 1,573 samples collected from 18 different body sites in healthy Americans. HPViewer detected 104 HPV types representing 4 HPV genera (Alpha, Beta, Gamma, Mu) [40] in 175 samples (Fig 6 and Table 3) in 16 of the 18 body sites (overall prevalence: 11.10%). Of the 104 HPV types detected, 84 types of HPV (81.73%) should not be detectable by the widest spectrum cervical HPV detection kit, The Linear Array® (37 types of HPV) [15]. Among the 104 HPV types, the top four most commonly detected types were HPV51,17,18, 89 while 10 types of high-risk (66.67%, 10/15) and 7 types of low-risk (58.33%, 7/12) HPV were detected among these healthy samples. The body site with the highest prevalence of HPV was left retroauricular crease (65.22%), followed by right retroauricular crease (54.84%), vaginal inroitus (50.00%), anterior nares (37.96%), and posterior fornix (32.41%) while HPV was not detectable in samples from palatine tonsil and blood. According to their profiles of HPV prevalence, tongue dorsum, buccal mucosa, supragingical plaque, ileal pouch, and stool were clustered as one group, and three vagina-related body sites—mid vagina, vaginal introitus, posterior fornix—were clustered together and three skin-related body sites—anterior nares, and left, right retroarticular crease—were clustered as another group (Fig 6; **Supplementary Table 3**). It indicated that HPV prevalence was associated with its habitat environment and supported previous studies that suggested gut, mouth, and skin have their own HPV diversity spectrums [41–44]. Co-occurrence of multiple HPV types in one sample were common with distinct patterns with respect to body sites (Fig 7; **Supplementary Table 4**). In the co-occurrence network, there were 89 types of HPV co-occurring with others at least once. This network shared some similarity with previous study [26]. Interestingly, HPV23-173 in skin, HPV54-89, and HPV39-51 in vagina were three most commonly observed co-occurrences (three times) and we did not find any co-occurrence relation shared between skin and vagina. These findings confirm that HPViewer is a broad range detection tool suitable for the evaluation of HPV presence beyond the female genital system.

**Table 3.**
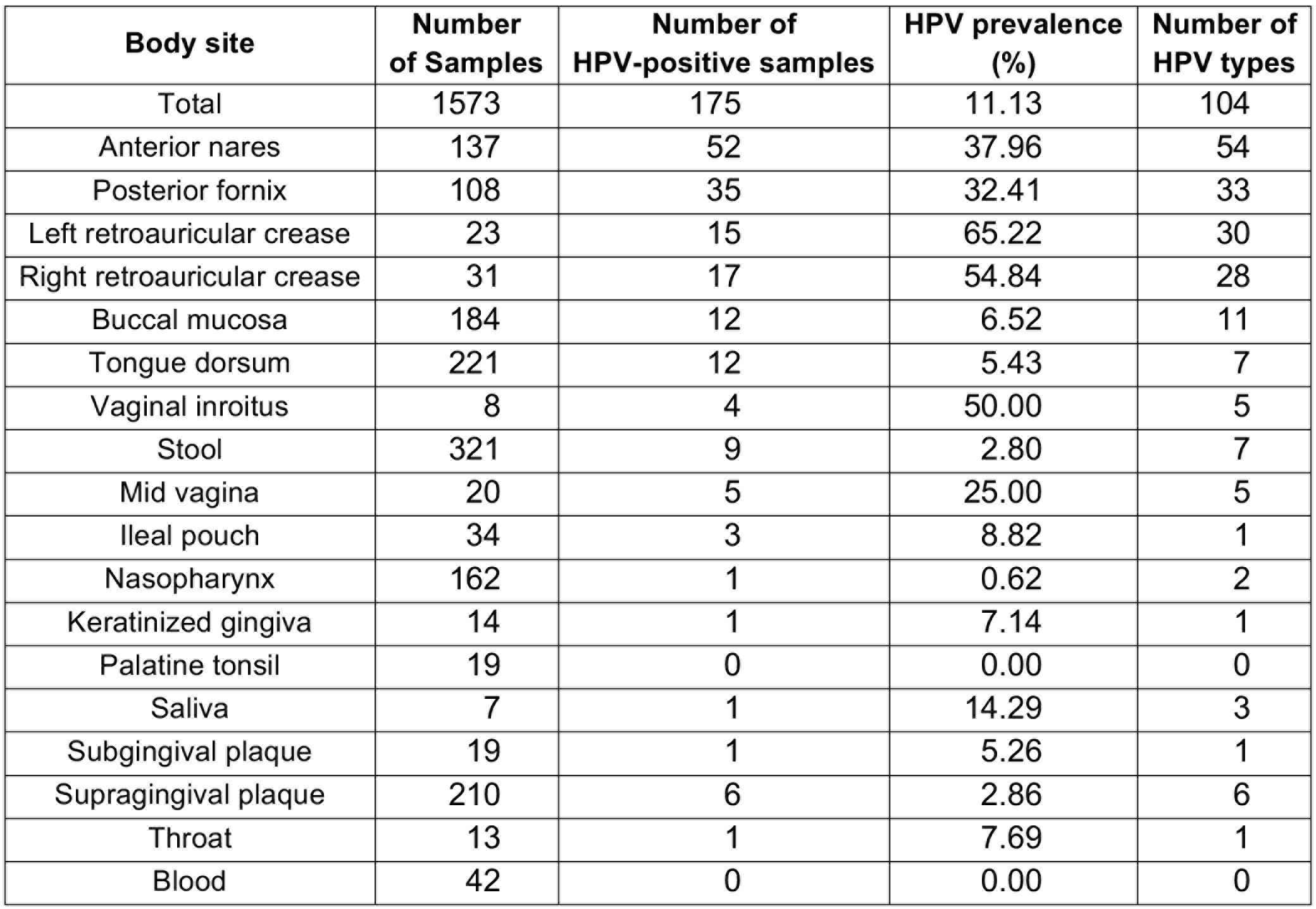
Summary of HPV-positive samples from the Human Microbiome Project

**Figure 6.**
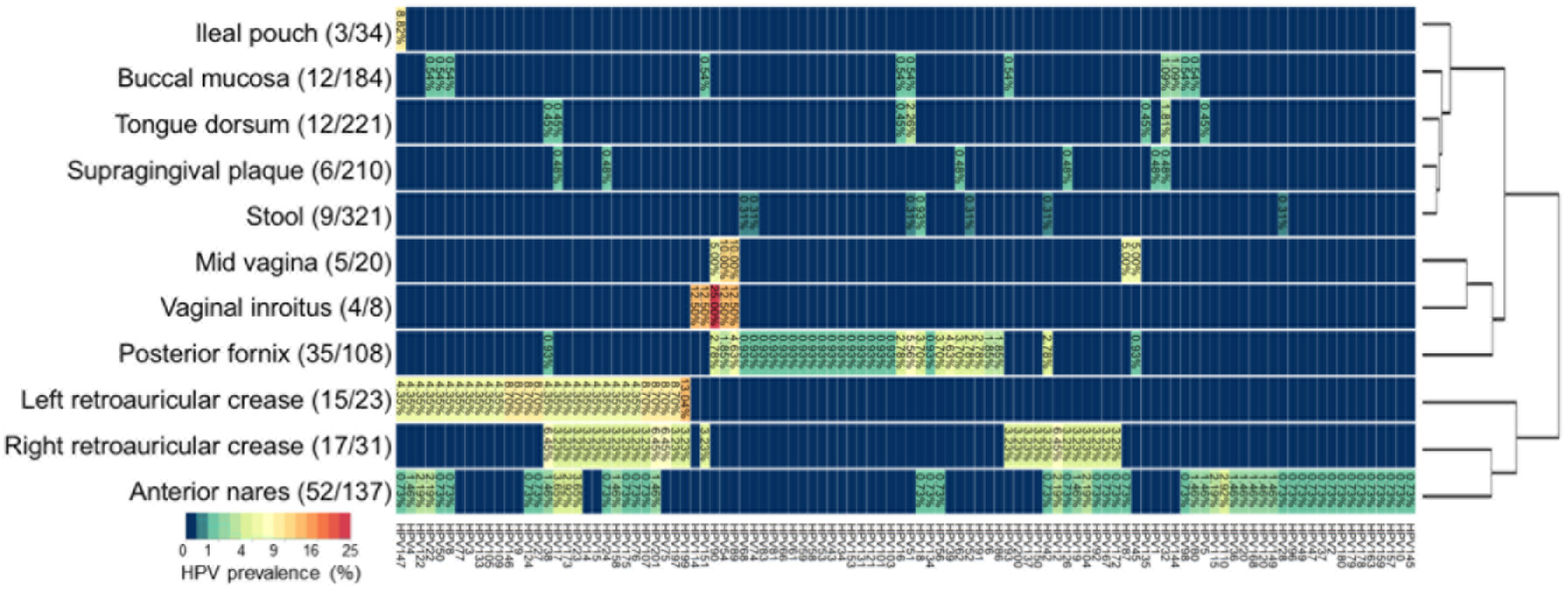
HPV prevalence summary of shotgun metagenomic data from HMP. 11 of 18 sites that were evaluated at least had two HPV positive samples. Sites are clustered vertically by their HPV prevalence pattern. The number in the parenthesis close to the body site label is the overall HPV positive sample / total samples and the number in the plot is the HPV prevalence for each HPV type in each body site.

**Figure 7.**
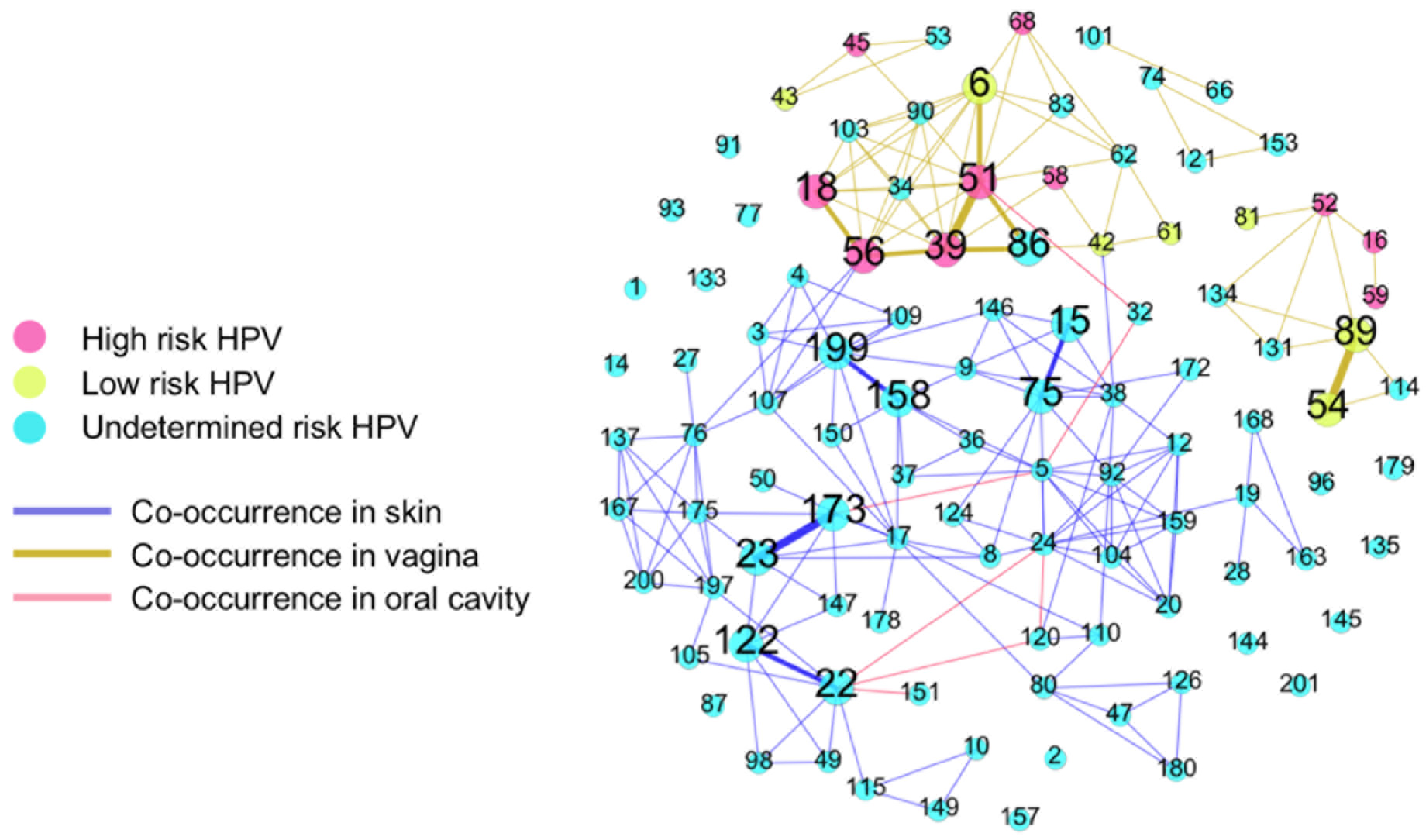
Co-occurrence graph of HPV in skin, vagina and oral cavity HMP samples. It consists of all 104 types of HPV. Each node represents one type of HPV and each edge represents the linked two nodes were found to co-existed. The thickness represents the frequency of co-occurrence in the range of 1 to 3. The nodes without any edges were not observed to have any co-occurrence. The skin includes anterior nares, left/right retroauricular crease; vagina includes mid vagina, posterior fornix, vaginal inroitus; oral cavity includes saliva, tongue dorsum, nasopharynx, and buccal mucosa. Most co-occurrence (edges) happened in skin or vagina and there were only six co-occurrence in the oral cavity.

## Discussion

HPV is an important human pathogen not only because it is the main cause of cervical, oropharyngeal and anal cancers but also because of the increasing evidence to suggest non-cervical HPV types might play an etiological role for cancers of many other body sites. Given the inadequacy of cervical HPV detection kits to cover all 210 HPV types, metagenomic shotgun sequencing has emerged as one of the most promising strategies for the detection of HPV in human samples. Now, we show that HPV not only shares a substantial amount of homologous sequences among different HPV types but also shares extensive simple repeats with human and some prokaryotes. With HPVDetector, a previously published software program specially designed for detecting HPV in metagenomic data, we found that the intra-HPV homologous sequences cause errors in HPV genotyping and the shared repeats of human or prokaryotes origin can be mistaken as HPV DNA, indicating a need to design a program for more accurate detection and genotyping of HPV.

A HPV type is defined if its major capsid L1 gene sequence is less than 90% similar to that of any other types [45]. In the present study, we found it is common that regions of one HPV type share high similarity (>90%) with other types despite their L1 genes share less than 90% similarity. In 28 types of HPV, the shared portions accounted greater than 50% of their genomes. These variations in similarity among HPV genomes make it difficult to create an operational threshold for accurate genotyping among HPV types using short reads generated from shotgun sequencing. Yet it is clinically important to accurately determine the type of HPV in each sample, since HPV types differ in their pathogenic properties. We created a type-specific database by removal of all regions that shared >90% similarity among HPV types from the HPV reference genomes (homology-mask). We used the type-specific database in HPViewer and demonstrated that the homology-mask mode of HPViewer can reduce misclassification of reads to less than 0.3%.

An ideal software program for detection and genotyping HPV from shotgun sequences should be both specific and sensitive. Some HPV types share simple repeats with the human genome and prokaryotic genomes. In the papillomatosis samples, HPVDetector misclassified TG repeats of human origin as HPV 71. VirusTAP takes two steps to ensure specificity. One is to filter out reads that are shared between HPV and non-HPV organisms and the second one which is also applied by Vipie, is to build up a large *de novo* assembled contigs to minimize the impact of local non-specific regions. This approach is demonstrated to be most specific among all programs evaluated. However, the high specificity is achieved at a cost of lower sensitivity due to the failure to assemble of contigs with sufficient length when a sample contains few HPV reads. In the papillomatosis study, VirusTAP failed to detect HPV 6 in tumor samples despite each sample containing hundreds of HPV 16 reads. Vipie is more sensitive than VirusTAP but unable to detect HPV in samples that contain less than 10 HPV reads. In contrast, HPVDetector is sensitive but less specific because of false positives from reads shared between HPV and human and prokaryotes or among HPV types.

HPViewer detects HPV by directly matching reads to HPV-specific reference genomes without *de novo* assembly. It achieved similar specificity to VirusTAP and Vipie with the threshold established by HPV type-specific PCR and higher sensitivity capable of detecting HPV in samples with as few as two HPV reads. The importance of detecting low HPV reads was exemplified in the study of recurrent respiratory papillomatosis. VirusTAP failed to recognize HPV infection in two (33%) of the six tumor samples despite several hundreds of HPV reads in the datasets, making it inadequate for diagnose of HPV infection in clinical samples. Vipie was unable to detect the presence of HPV 6 in two oral samples in which there were two HPV reads in each sample. Because these four reads belonged to the same strains in the corresponding papilloma tumor samples, failure to detect them might underestimate the potential transmissibility of HPV from recurrent respiratory papillomatosis through the oral route.

## Conclusions

In summary, HPViewer is a new tool designed for broad range detection and genotyping of HPV in shotgun sequencing data from human samples. It has high sensitivity by directly detecting HPV from raw sequence reads. It eliminates false positives by masking simple repeats in the reference HPV genomes shared by human and bacteria and reduces mistyping of HPV reads by masking homologous sequences shared among different HPV types. To optimize the trade-off between sensitivity and specificity, the hybrid mode of HPViewer integrates these two kinds of masked HPV genomes using the pair-wise homology distance matrix.

What is more, it uses the least space for data storage and provides faster time for analysis of HPV in a sample compared with other software programs available. HPViewer also has a built-in function to calculate HPV genome coverage. HPViewer is implemented with python and operates in the Linux environment so it can easily be used to process large numbers of samples. It produces a table containing HPV types detected, the number of matching reads and their depth of coverage on reference HPV genomes, and a bam file containing short reads aligned to HPV genomes, which can be visualized in the Integrative Genomics Viewer [46]. With the rapidly decreasing cost of shotgun sequencing, metagenomics has emerged as one of the most effective strategies for the detection of HPV in clinical samples. HPViewer is a sensitive and specific tool for use in the analysis of HPV infection.

## Methods

### Two HPV genome databases in HPViewer

We downloaded all 182 HPV reference genomes from PaVE for this study. Bowtie2 (version 2.2.7) is the alignment tool utilized in this study [47]. All metagenomic reads were aligned to our customized HPV databases through bowtie2 in the end-to-end, sensitive mode.

We created two local HPV databases with two different masking strategies, repeat-mask and homology-mask. For the repeat-mask database, we used RepeatMasker to replace the low complexity and simple repeats regions of all HPV genomes with “N”. For the homology-mask database which was inspired by Metaphlan [48], we created a type-specific HPV database by masking homologous sequences shared among different HPV types, and then further masked the repeats using RepeatMasker (**Supplementary Fig 1**). There were three steps for the construction of homology-mask database. First, all 100 bp DNA fragments from each complete HPV genome generated by EMBOSS [49], were aligned to all other types of HPV with a 90% identity threshold by bowtie2 (bowtie2 parameters: -a --score-min L,0.6,0.6). Then we masked the matching regions on the genomes (**Supplementary Fig 1**). Finally, after all homologous regions were masked, RepeatMasker was also applied for all processed HPV genomes to mask low complexity and simple repeats regions (**Supplementary Fig 1**). For repeat-mask and homology-mask databases, the length of HPV genomes was not changed and only some fragments were replaced as ‘N’, and we called non-N sequences of HPV genome as the effective genome. The distribution of effective genome size of original HPV, repeat-mask, and homology-mask was generated by the R package, ggplot2 [50].

We validated the repeat-mask database by BLASTn against genomes of human (GCRh38) and prokaryotes (Prokaryotic RefSeq 112 reference genomes and 1,669 representative genomes) (https://www.ncbi.nlm.nih.gov/genome/browse/reference/), and no matches were found with identity > 90% over an alignment region > 50 bp. The circos plot of shared sequences between HPV and human, prokaryotes was generated by Circos 0.69 [51].

### Construction of the homology tree among 182 types of HPV and the hybrid-mask of HPViewer

In order to explore the sequence similarity among HPV types, we selected from each genome only the 100-bp genome fragments with >90% identity to two or more HPV types from each HPV type, all other bases in the genomes were masked as N (**Supplementary Fig 1**). There were 29 types of HPV without any 100-bp regions that matched other types. The selected portions from the remaining 153 HPV genomes were multiple-aligned with MUSCLE 3.8.31 [52].The pairwise distance matrix was calculated by MEGA7 [53] and the maximum likelihood tree was built with RAxML 8.2.9 under a GTRCAT substitution model with 1,000 bootstrapping replicates [54]. The homology tree with a midpoint root was visualized by FigTree v1.4.3 [55] (**Supplementary Fig 2**).

For the hybrid-mode of HPViewer, first, the repeat-mask mode is used to identify all HPV types in a sample. We set the threshold of detection of one HPV type in a sample as two different aligned reads covering at least 150 bases of a single HPV type reference genome, (**Supplementary Fig 4**). We used SAMtools depth [56] to obtain the coverage for each position of mapped HPV genomes. When the length of the covered positions of the mapped reads on a single HPV type is smaller than 150, we discard that HPV type as false positive. When the covered length is above 150 bp, we considered it as detected.

When only a single HPV type is detected in the sample data file, there is no chance for false positives from other HPV types, so it is considered as a true positive. When multiple types of HPV are detected in a sample, the HPV types are checked if they are close to each other (the homology distance < 0.35) using the pair-wise homology distance matrix. Distantly related HPV types are reported directly in the HPV profile. The closely related HPV types are required to be re-tested, thus HPV reads generated from repeat-mode output bam file by BEDtools [57] are re-aligned to the homology-mask database. Only similar HPV types detected by homology-mask mode are also added into the HPV profile.

### Simulation of HPV shotgun sequencing data with Grinder

Simulated HPV samples used in our model evaluation were produced by Grinder [58] and each sample contains one of 143 types of HPV which are detectable by HPVDetector. For each type of HPV, we generated 100 samples with seven different levels of HPV reads mimicking different sequencing depth: 2, 5, 10, 50, 100, 500, and 1000. In total, there were 100,100 simulated samples (143*100*7). Reads 100bp long were sampled from the selected genomes adding 5% mutations to increase diversity.

### Detection and genotyping of HPV in patients with recurrent laryngeal papillomatosis using HPViewer

Following approval from the Institutional Review Board at the New York University School of Medicine (study number S13-00119), six patients with pathology-confirmed recurrent respiratory papillomatosis were identified from large pool of patients participating in a large-scale, longitudinal study. Tumor tissue was endoscopically removed, fixed in formalin, and embedded in paraffin. The diagnosis of recurrent respiratory papillomatosis was made by histopathological examination of the tumor tissue. To extract DNA, the paraffin-embedded tissue was cut into 20 micron-thick sections. Total genomic DNA was extracted from the unstained tissue sections using BiOstic FFPE Tissue DNA Isolation Kit (Mo Bio Carlsbad, CA).

Oral rinse samples were collected from the same six patients according to the National Health And Nutrition Examination Survey (NHANES) protocol [59]. Briefly, subjects were instructed to swish 5mL of Scope® mouthwash without gargling for one minute. The oral wash samples were then sealed and stored for no more than one week at 4°C prior to DNA extraction. For DNA extraction, the samples were spun for 10 minutes at 3200g. DNA in the cell-free supernatant was precipitated with the isopropanol/glycogen solution and pelleted for 10 minutes at 2000g, as previously described [59]. The pellet was resuspended with 200 μL DNA Hydration Solution (Qiagen).

To evaluate the sensitivity and specificity of HPViewer, we determined the true HPV compositions in these papilloma and oral wash samples. Since only HPV types 6 and 11 have been previously observed in laryngeal papilloma [37], we conducted standard PCR for HPV6 and 11 using the primers from Tucker et al. (2001) [60] on these 12 samples. We found HPV6 in 4 tumor tissues and 2 matched oral wash samples and HPV11 in 2 tumor tissues. Additional PCR with lower annealing temperature confirmed that HPV6 was present in samples 3W and 7W and that both 3T and 3W were negative for HPV11 (**Supplementary Fig 5**).

In the original Bowtie2 screening of these samples on an unmasked HPV database, small numbers of reads matching HPV71 were found in all six oral samples and three tumor samples, as well as HPV19 and 82 in some samples. Inspection of these reads revealed sequences such as TG repeats (**Supplementary Fig 3)** which matched to TG repeats in the genome of HPV71. After masking the HPV database with RepeatMasker, no reads matching HPV19, 71, or 82 were found.

HPViewer identified just 2 reads of HPV6 in oral wash samples 3W and 7W. These samples were confirmed as HPV6 positive by PCR. Inspection of the sequence of these reads revealed that they contained the same polymorphisms found in the much larger number of reads in matched tumor samples from the same patients, suggesting a low level of release of HPV from the papilloma into the oral cavity. HPViewer detected just 1 read of HPV11 in sample 3W, but HPV11 was not detected by the PCR in 3W or 3T. Consequently, we have set the detection threshold for HPViewer at 2 different reads per sample for a single HPV type.

### Metagenomic data from Human Microbiome Project

We downloaded 1,573 shotgun sequencing metagenomic data sets from Human Microbiome Project (https://hmpdacc.org/hmp/) (**Supplementary Table 5**). The HMP samples (with human data previously removed) were obtained from 18 body sites, including anterior nares, attached keratinized gingiva, blood, buccal mucosa, ileal pouch, left retroauricular crease, mid vagina, nasopharynx, palatine tonsils, posterior fornix, right retroauricular crease, saliva, stool, subgingival plaque, supragingival plaque, throat, tongue dorsum, and vaginal introitus (**Supplementary Table 3**). The heatmap of HPV prevalence for different body sites were produced by R package, gplot [61]. The co-occurrence of HPV for three body sites were generated by Gephi 0.9.1 [62].

## Declarations

### Ethics approval and consent to participate

Oral Cavity Human Papilloma Virus Reservoirs in RRP” with study number: S13-00119.

### Consent for publication

Not applicable.

### Availability of data and materials

HPViewer is freely available as a python package at https://github.com/yuhanH/HPViewer under the terms of the GNU General Public License (version 3) as published by the Free Software Foundation for academic use. Commercial users should contact Dr. Pei at. All RefSeq reference and representative genome sequences used in our analyses are publically available online via NCBI and detailed information can be found in the Table S2. All 1,573 HMP metagenomic dataset could be found in the HMP website (https://hmpdacc.org/hmp/) and their SRS ID were listed in the Table S5. The prevalence and co-occurrence data of HMP metagenomic samples can be found in the Table S3 and Table S4.

### Authors’ contributions

Each author has contributed substantially to the work, and has edited and approved this manuscript.

## Acknowledgements

We thank the Applied Bioinformatics Laboratories at NYU School of Medicine for providing bioinformatics support and assisting with data analysis. This work utilized computing resources at the High Performance Computing Facility at NYU Langone Medical Center. We also thank the Genome Technology Center for expert sequencing. This shared resource is partially supported by a Cancer Center Support Grant, P30CA016087, at the Laura and Isaac Perlmutter Cancer Center. ZP is staff physician at the Department of Veterans Affairs New York Harbor Healthcare System. The content is the sole responsibility of the authors and does not necessarily represent the official views of the National Institutes of Health, the U.S. Department of Veterans Affairs or the United States Government.

